# Strigolactone mediates moso bamboo root response to phosphate stress

**DOI:** 10.1101/2022.07.21.501044

**Authors:** Qian Wang, Ping Yang, Muhammad Asim, Renyi Gui, Mingbing Zhou

## Abstract

Moso bamboo (*Phyllostachys edulis*) grows in soils with widespread phosphate (Pi) deficiency, resulting in low shoot yield. While it also suffers from high Pi due to heavy fertilization, which causes the degradation of bamboo forest. A novel plant hormone, strigolactone (SL), plays a key role in root growth and development under Pi stress, but its regulatory mechanism has not been systematically reported. Our study investigated SL-mediated mechanism in response to Pi stress in moso bamboo. We compared the root growth under low, sufficient and high Pi, and analyzed the temporary trends of gene expression in primary root tip region and lateral root primordium zone. The effects of SL-analog (GR24) and SL-inhibitor (TIS108) on root architecture at low and high Pi were evaluated. SL biosynthesis and signaling are the main pathways of root response to Pi stress. With the decrease of Pi level, 5-deoxystrigol and strigol in the root exudates increase significantly. Under low Pi, SL is indispensable for the maintenance of root cell morphology, and promotes primary root elongation and reduces lateral root formation by upregulating the expression of phosphorous starvation response genes and downregulating the expression of abscisic acid response genes. The absence of SL at high Pi releases the inhibition of ethylene responsive genes’ expression to inhibit root elongation and promote branching. In general, SL mediates the response of bamboo roots to Pi stress by regulating its biosynthesis and signal transduction and influencing other hormone pathways.

## 1. Introduction

Bamboo is one of the world’s most valuable non-timber forest products, with an international trade value of USD 3.054 billion in 2019 (INBAR, 2021). Unlike other arbor and shrub species, bamboo has a short stem lifespan, a long flowering cycle and propagates vegetatively with a robust reproductive rhizome. Moso is a large woody bamboo species with the highest ecological, economic and cultural value, accounting for more than 70% of the growing area of bamboo. However, moso is widely distributed in the mountainous areas of southern China where the soil is generally phosphate (Pi) deficient, resulting in low yield of bamboo shoots, but it grows well with high-developed root system. Under low Pi condition, the phosphorus acquisition efficiency (PAE) in bamboo can be improved by root architecture alteration, organic acids secretion and interaction with soil microorganisms (Ruangsanka, 2014). On the contrary, Pi fertilizers are commonly used to promote shoots production in intensive-managed bamboo forests, but excessive fertilization can also lead to soil acidification (Gui et al., 2013), imbalanced mineral element ratios in soil and bamboo, aluminum toxicity (Gui et al., 2011), and ultimately bamboo forest degradation (Gui et al., 2018). Therefore, to improve the cultivation efficiency of bamboo forest and protect the soil ecological environment, it is particularly important to develop technical measures to improve PAE and excavate bamboo varieties resistant to Pi deficiency.

Phosphorus, acquired in the form of Pi, is one of the primary macronutrients for plants but is least available in the soil (Czarnecki et al., 2013). Fine-tuning of root system architecture (RSA) is the most obvious and effective response of plants to Pi stress. For example, Arabidopsis has shorter primary root and more lateral roots under Pi deficiency (Mayzlish-Gati et al., 2012), primary root length increases significantly in rice (Sun et al., 2014), and total root length of wheat has a significant positive correlation with Pi uptake (Newman, 1973). Plants also have their adaptation mechanism under Pi toxic conditions with polymorphism in different plants. Local reapplication of Pi inhibits primary root elongation and promotes lateral root formation of Arabidopsis (Linkohr et al., 2002). In rice, high Pi such as 120 kg/ha inhibits root elongation and root number (Yang et al., 2021).

Interestingly, the adaptive changes of plants under Pi stress are jointly regulated by multiple plant hormones, among which strigolactone (SL) plays a key role (Rasmussen et al., 2013). Pi deficiency stimulates SL biosynthesis in the roots, and SL can improve PAE by modulating RSA. SL is also transported through xylem to inhibit branching, a means of reducing Pi utilization (Yoneyama et al., 2007; Czarnecki et al., 2013). Effects of SL on shoots are similar in most plants -it interacts with auxin to inhibit branching, but reports on root development in different plants are inconsistent. Under low Pi, SL inhibits root elongation and promotes lateral root formation in Arabidopsis (Ruyter-Spira et al., 2011). However, SL promotes primary root elongation and inhibits lateral root formation in rice (Sun et al., 2014). Under sufficient Pi, application of SL analog GR24 has no effect on primary root elongation, but promotes primary root growth in rice; and Pi starvation responses (PSRs) like inhibition of lateral root formation are induced both in Arabidopsis and rice. SL may manipulate the auxin pathway to regulate roots under low Pi in more than one way: inhibiting auxin transport by consuming PIN family on the plasma membrane and increasing auxin perception by promoting *TRANSPORT INHIBITOR RESPONSE1* (TIR1) transcription (Shinohara et al., 2013; Pandya-Kumar et al., 2014; Koltai, 2015). SL also enhances Pi uptake by fine-tuning the activity of phosphorous starvation inducing genes like *PHOSPHATE OVER-ACCUMULATOR 2* (*PHO2*) and regulating of root hair elongation and lateral root formation (Kumar et al., 2015; de Souza Campos et al., 2019). These processes are dependent on MORE AXILLARY GROWTH 2 (MAX2), a key signaling component in SL pathway (Mayzlish-Gati et al., 2012). The exact role of SL in the regulation of root structure is still unknown currently, and there is no unified evidence of downstream signal events of SL-mediated root development responding to Pi stress.

In this research, we observed the root morphological changes, cell microstructure characteristics and endogenous hormone changes of moso bamboo under low, sufficient and high Pi. Transcriptome sequencing was carried out in primary root tip and LRP zone of moso bamboo roots to analyze the dynamic changes of gene expression under Pi stress and explore the regulatory mechanism of SL in root development. The effects of exogenous SL analog and inhibitor on bamboo root architecture verified the molecular mechanism of SL participating in bamboo root response to Pi stress.

## 2. Materials and methods

### 2.1 Plant materials and growth condition

Moso bamboo seeds were soaked in deionized water in a dark incubator at 30 °C for two days and then transferred on a net floating in a 0.5 mM CaCl_2_ solution in a 1L plastic container. After 5 days, seedlings were removed from seeds and transferred to 1/2 Kimura B nutrient solution (pH5.6) for 7 days. The solution contained the following nutrients (mM): MgSO_4_ (0.28), (NH_4_)_2_SO_4_ (0.18), Ca(NO_3_)_2_ (0.18), KNO_3_ (0.09), and KH_2_PO_4_ (0.09), Fe-EDTA (20), H_3_BO_3_ (3), MnCl_2_ (0.5), CuSO_4_ (0.2), ZnSO_4_ (0.4), and (NH_4_)_6_Mo_7_O_24_ (1) (Hu et al., 2018).

1. The 16-day-old seedlings were then transferred to nutrient solutions with different Pi concentrations for 4 weeks. KH_2_PO_4_ was replaced with 0, 1, 2, 50, 100, 500, 1000, 2000 μM NaH_2_PO_4_, while K_2_SO_4_ (0.09mM) was used to maintain sufficient supply of potassium. Root parameters were analyzed to determine low, sufficient and high Pi concentrations.
2. The 16-day-old seedlings were then transferred to nutrient solutions with sufficient Pi (100 μM) containing 0.01, 0.1, 1, 2 μM GR24 or TIS108 for 2 weeks to determine the effective GR24/TIS108 concentration (Ito et al., 2013; Oláh et al., 2020).
3. The seedlings were then transferred to nutrient solutions containing low Pi (1μM), sufficient Pi (100 μM) or high Pi (1000 μM) without or with 1 μM GR24 or TIS108 for 2 weeks.

All solution was renewed every two days and all experiments were conducted with at least three biological replicates.

### 2.2 Root morphological analysis

Primary root length, lateral root number and average diameter were measured by LA-S Software (Hangzhou Wanshen Detection Technology Co., Ltd. China). 0.3-0.5cm of primary root tip and 1.5-3cm of LRP zone were collected and cut into 40µm by Leica VT1200S fully automated vibrating blade microtome (Leica, Germany). The anatomical characteristics of primary root apical meristem and lateral root primordia were observed by Zeiss Axio Imager. A2 (Carl Zeiss Jena, Germany).

### 2.3 Quantification of total phosphorous content

At the end of the experiment, shoots and roots were harvested and dried at 70℃ to measure the biomass. Total phosphorus content was determined by vanado-molybdate yellow method (Yoneyama et al., 2012).

### 2.4 Extraction and quantification of SL

Root exudate was collected after 96h, 192h and 336h after treatment with different Pi concentrations. The nutrient solution was filtered with activated carbon and then fully mixed with 2 mL acetone, which was passed through the pretreated HLB solid-phase extraction column, eluted with 2 mL 100% acetone, blow-dried acetone with nitrogen, and redissolved with 200 uL acetonitrile. Liquid mass spectrometry (HPLC-MS/MS) was used for detection (Rial et al., 2019).

### 2.5 Total RNA extraction and library construction

At 24h, 48h, 96h, and 336h of low Pi, sufficient Pi and high Pi treatment, 0.5 cm primary root tips (including meristem zone and elongation zone) were collected as described previously (Zhang et al., 2019). At 48h, 96h, 192h and 336h of low Pi, sufficient Pi and high Pi, 1.5 cm LRP zone of primary root was collected as described previously (Li et al., 2012). Total RNA was extracted using RNAprep pure plant plus kit (Tiangen, Beijing, China) for cDNA library construction. Sequencing was performed using the Illumina HiSeq 2500 (Illumina Inc., New York, USA) platform based on sequencing by synthesis (SBS) technology to generate large amounts of raw data. Then adapter was removed and low-quality reads were filtered to ensure high quality clean data and accuracy.

### 2.6 Transcriptome data analysis

Cluster and ggplot2 in R language were used for principal component analysis (PCA). Cufflinks was used for quantitative analysis of transcriptome expression, where the amount of expression was measured by fragments per kilobase million (FPKM). DEGs between samples were screened by fold change (FC) using edgeR and DESeq2 with |log2 Ratio|≥2 and FDR≤0.005. Weighted correlation network analysis (WGCNA) was conducted to build the co-expression network of all genes by using pickSoftThreshold and powerEstimate to calculate the soft threshold and the optimal weight value respectively. Modules were preliminarily identified by dynamic shearing tree method, and the co-expression network was constructed by merging modules similar to ME (Module eigengene E). Genes with the highest connectivity in the module were selected as the core hub genes of the module through Pearson correlation method (Cor>0.5, *P*<0.05), and co-expression network was visualized using Cytoscape V3.7.0. Gene function was annotated and enriched using Gene Ontology (http://geneontology.org), KEGG (http://www.kegg.jp/kegg/pathway.html), BambooGDB (http://bamboo.bamboogdb.org) and Uniprot protein database (https://www.uniprot.org). DEGs expression trends were displayed using tools on BMKCloud platform (https://international.biocloud.net).

### 2.7 Real-time RT-PCR of candidate genes

Twenty-three DEGs were selected for qRT-PCR verification. The length of the amplified sequence ranged from 100 bp to 300 bp, and primer sequences are shown in Table S4. 1μg of DNAase-treated total RNA was used for reverse transcription with reverse transcriptase (Yeasen Biotechnology (Shanghai), Co., Ltd. China) in a final reaction mixture of 20 ul. Two-step qRT-PCR was performed on Bio-RadCFX96^TM^ with SYBR Green. The reaction process includes initial denaturation at 95℃ for 5 min, 40 cycles at 95℃ for 10s and 60℃ for 30s, and a final melting curve stage at 65-95℃ for 5s. The relative gene expression level was calculated using 2^-ΔΔCq^ method according to previous studies (Abdallah et al., 2016) with *NTB (NUCLEOTIDE TRACT - BINDING PROTEIN*, Genbank accession: gi|242381788) gene as an internal control. Four biological replicates were set for all reactions, and the specificity of primers was verified by agarose gel electrophoresis and dissolution curve analysis.

## 3. Results

### 3.1 Determination of low Pi, sufficient Pi and high Pi level

After 4 weeks of treatment, root biomass of moso seedlings gradually increased with the highest under 0-1 μM and the lowest under 2000 μM as the decrease of Pi concentration, while shoot biomass increased gradually with the increase of Pi concentration and reached the highest at 100 μM and then decreased slightly (Fig.S1A). This suggests that 100 μM provides sufficient Pi for the initial growth of moso seedlings. When Pi concentration was above 500 μM, the leaves gradually turned yellow, grew poorly, and died after 3-4 months of treatment, indicating that Pi concentrations above 500 μM probably have toxic effect on the seedlings. Root-shoot ratio increased and total phosphorous content decreased with the decrease of Pi concentration (Fig.S1BC), indicating that Pi can change the proportion of biomass allocation in different tissues. We also analyzed the effect of Pi levels on root parameters. With the decrease of Pi concentration, primary root length, average lateral root length and average root diameter increased gradually (Fig. 1A, Fig.S1EF). In contrast, total root length and lateral root density decreased gradually (Fig.S1D, Fig. 1B), with the most significant differences between every two Pi levels in primary root length and lateral root density. Compared with under 100 μM, primary root length increased and lateral root density decreased significantly under 1 μM, while primary root length decreased and lateral root density increased significantly under 1000 μM, so 1, 100 and 1000 μM were selected as low Pi (LP), sufficient Pi (CK) and high Pi (HP) levels (Fig1. AB, Fig.S1GH).

**Figure 1.**
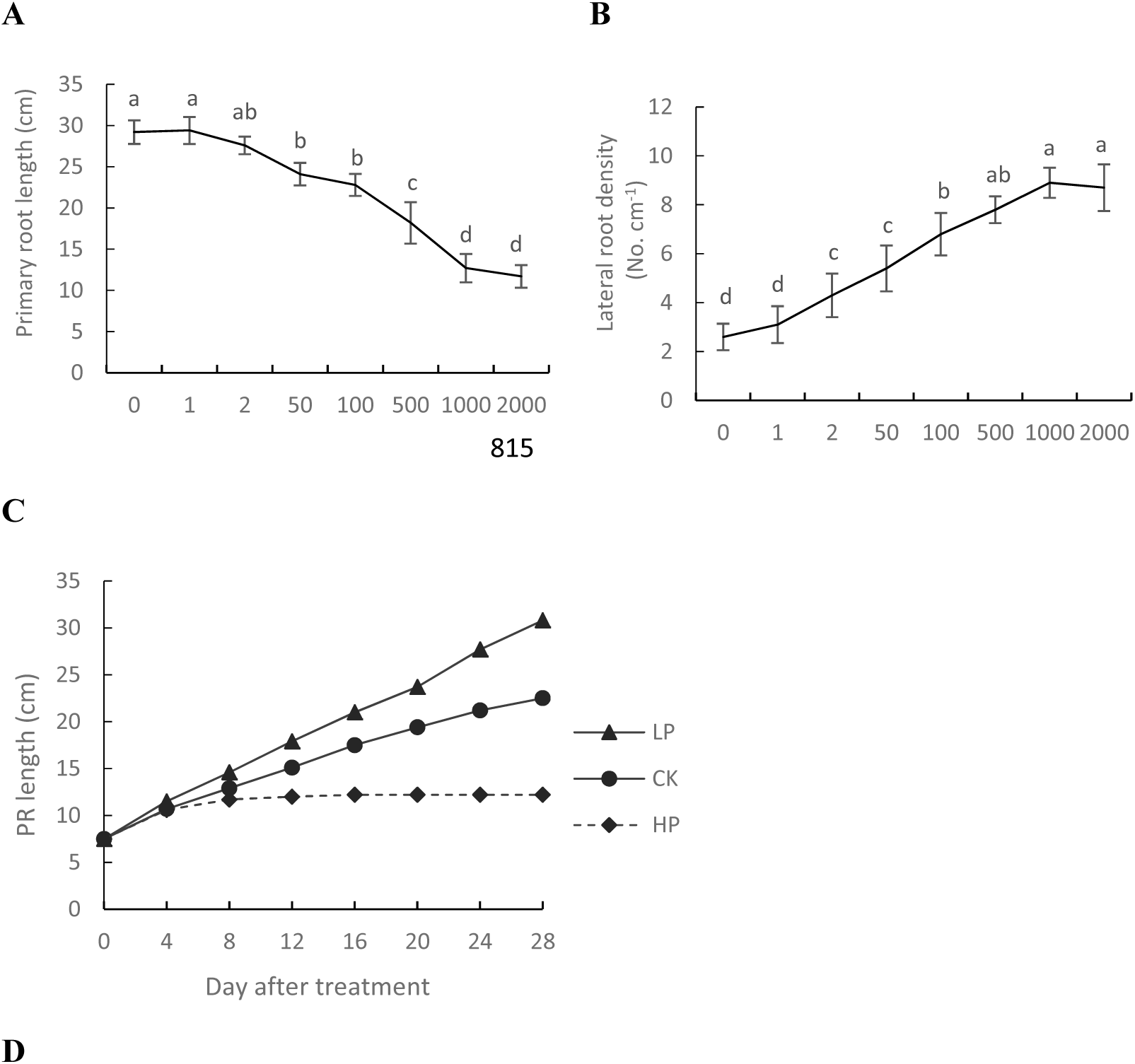
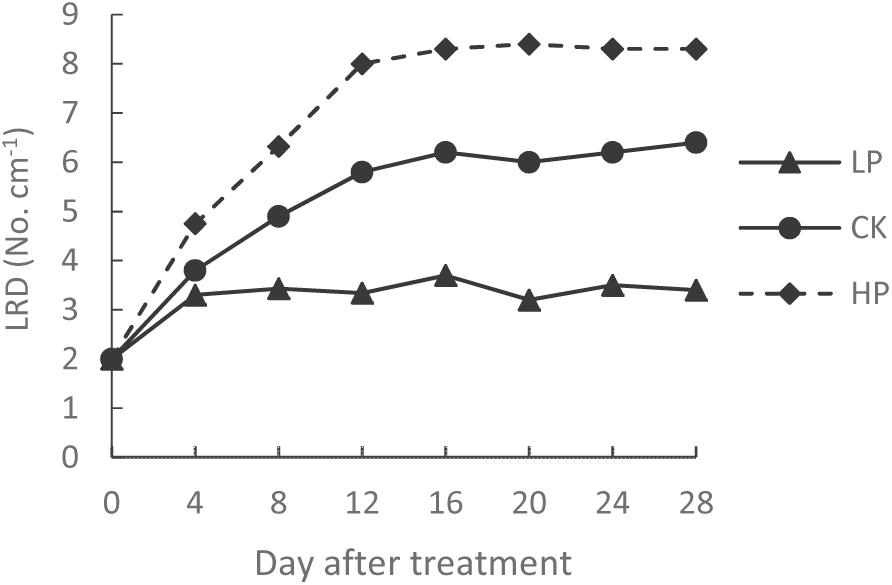
Effect of Pi availability on primary root length and lateral root density of moso bamboo. (A)(C) Primary root length and (B)(D) lateral root density of moso bamboo seedlings grown for 4 weeks in hydroponic solution with different Pi concentrations.

### 3.2 Dynamic change of primary root length and lateral root density of moso bamboo under Pi stress

In order to determine the specific time of root response to Pi stress, the effects of low Pi, sufficient Pi and high Pi on primary root length and lateral root density of moso bamboo were observed every 4 days (Fig. 1CD). Primary root length under low Pi was significantly lower than that under high Pi on day 4 (96h after treatment), and significant differences appeared among the three Pi levels on day 16 (Fig. S2A). Lateral root density under high Pi was significantly higher than under low Pi at day 8 (192h after treatment), and there were significant differences among all three treatments at day 16 (Fig. S2B). The primary root tip stopped growing and became browning at the 16th day under high Pi. To avoid root tip vitality loss when sampling, primary root length was monitored daily after 12 days of treatment. Significant differences were found both in primary root length and lateral root density among low, sufficient and high Pi at the 14th day (336h after treatment) (Fig. S2CD). Therefore, we harvested primary root tips at 24h, 48h, 96h and 336h and LRP zones at 48h, 96h, 192h and 336h after treatment for RNA extraction.

### 3.3 The transcriptome of LRP zone and primary root tip responds to Pi stress

To better understand root growth in response to Pi stress, transcriptome sequencing experiments were performed under the same experimental conditions. In LRP zone, a total of 2297 genes were significantly differentially expressed among the three Pi treatments at four time points (Table 1). GO enrichment analysis of overlapping genes at any two time points showed that the biological processes responding to Pi stress contained 645 terms, accounting for 63% of all terms enriched (Table S1). Twenty biological processes were significantly enriched (Table 2), and most of them belongs to “cellular process”, “metabolic process”, “biological regulation” and “response to stimulus”, which involved both local and systematical signal transduction. In 15 biological processes such as “strigolactone biosynthesis process”, “cellular response to phosphate starvation” and “phosphate ion transport”, the average fold change (AFC) of DEGs between low and sufficient Pi gradually increased with time. It peaked at 336h, and the DEGs between low and high Pi had the same trend with a higher AFC (Table 2). This suggests that the DEGs in these biological processes were gradually significantly up-regulated under low Pi, and there was an interaction between hormone pathways such as strigolactone biosynthesis process and response to phosphate starvation. In other 5 biological processes, such as “cell response to strigolactone” and “negative regulation of cytokinin activation signaling pathway”, the AFC of DEGs were significantly down-regulated at 96h under low Pi, indicating a significant interaction between Pi signal perception and hormone signal transduction in the LRP zone of moso bamboo.

**Table 1.**
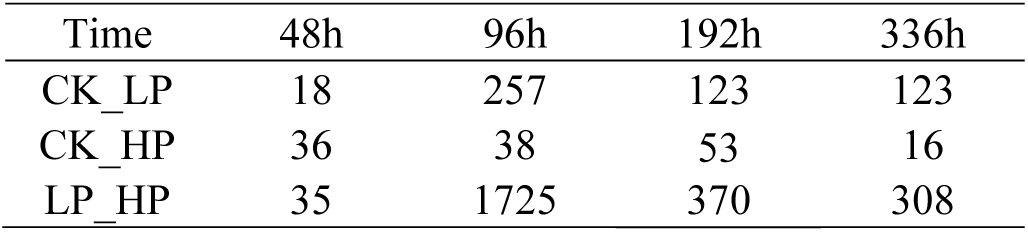
Differentially expressed gene (DEG) number in LRP zone in response to Pi stress

| Time | 48h | 96h | 192h | 336h |
| --- | --- | --- | --- | --- |
| CK_LP | 18 | 257 | 123 | 123 |
| CK_HP | 36 | 38 | 53 | 16 |
| LP_HP | 35 | 1725 | 370 | 308 |

**Table 2.**
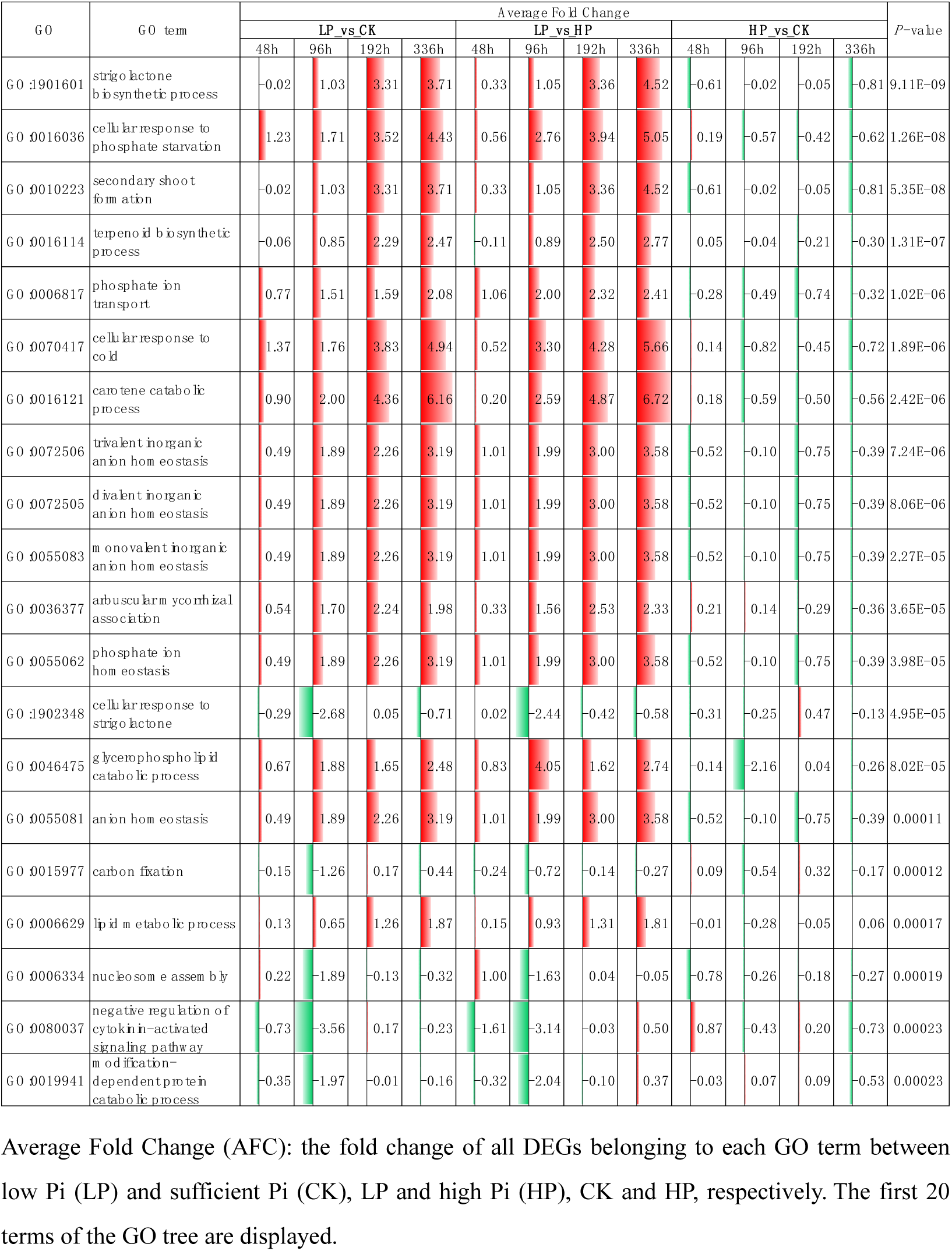
Significant enriched biological processes in LRP zone in response to Pi stress

In primary root tips, 5092 genes were significantly differentially expressed among the three Pi treatments at four time points (Table 3), with the most DEGs occurring at 96h and 336h, subsequently significant differences appeared in primary root length (Fig.S2AC). We also conducted GO enrichment analysis of overlapping genes at any two time points, and the biological processes responding to Pi stress contained 1833 terms, accounting for 67.9% of all terms enriched (Table S2). A set of 20 biological processes were significantly enriched (Table 2), and most of them also belongs to “metabolic process”, “biological regulation”, “response to stimulus” and “cellular process”. In “cellular response to phosphate starvation”, “divalent inorganic anion homeostasis”, “trivalent inorganic anion homeostasis”, “monovalent inorganic anion homeostasis” and “chemical homeostasis”, the AFC was significantly gradually up-regulated over time under low Pi and slightly down-regulated under high Pi comparing with sufficient Pi. Similar to the LRP zone, DEGs between low and high Pi has the same expression trend as DEGs between low and sufficient Pi but with a higher AFC. In 6 biological processes such as “negative regulation of cytokinin-activated signaling pathway”, AFC between low and sufficient Pi showed down-regulation with the highest AFC at 24h but then decreased gradually, indicating the rapid response and adaptability of root tip to low Pi. While the AFC between high and sufficient Pi were significantly upregulated at 96h, the time when primary root length was significantly decreased under high Pi comparied with under low Pi, indicating these processes may be involved with the root elongation. In the remaining 9 biological processes such as “organonitrogen compound catabolic process”, |AFC| among three Pi treatments at any time were all less than 1.5. In summary, many DEGs were found in LRP zone and primary root tips in response to Pi stress and involved many pathways. Most of them occurred between low Pi and sufficient Pi/high Pi. The AFC increased gradually over time, similar to primary root length and lateral root density, which became more significant with root development.

**Table 3.**
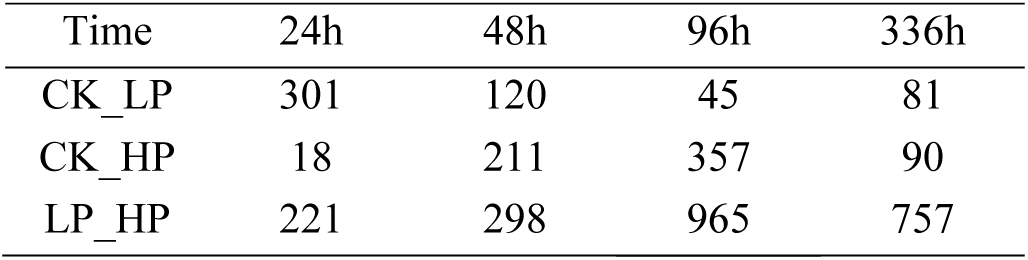
DEG number in primary root tips in response to Pi stress

**Table 4.**
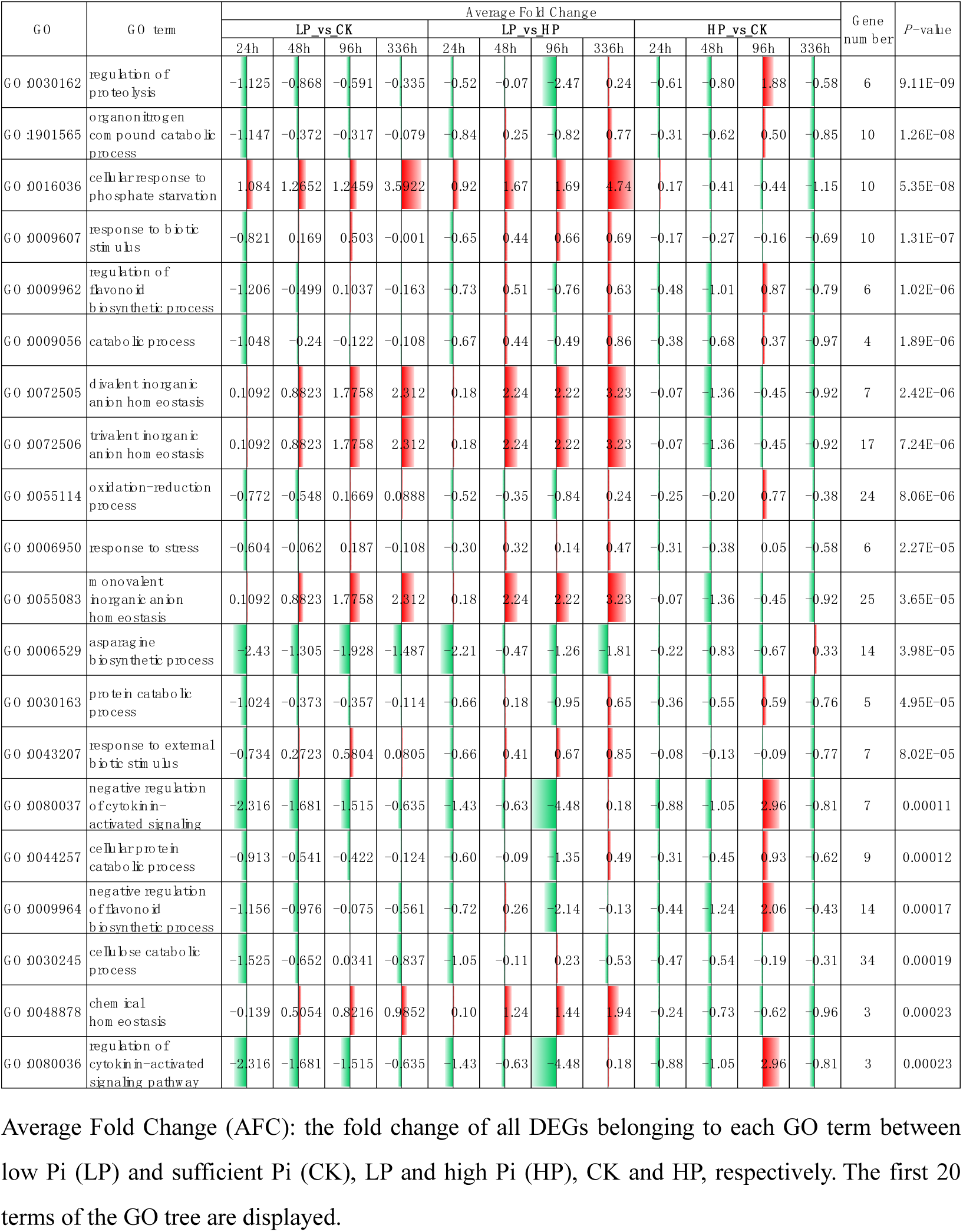
Significant enriched biological processes in primary root tips in response to Pi stress

### 3.4 WGCNA analysis

In order to investigate the correspondence between traits and genes and to screen out the most critical pathway of bamboo root response to Pi stress, WGCNA package in R was used to analyze gene expression matrix of all samples from primary root tips and LRP zone. Gene number in 14 modules varies greatly, with the least mediumpurple2 module containing only 47 genes and the most lightcyan1 module containing 1425 genes (Fig.S3AB). Module salmon, oranged3, mediumpurple2, skyblue3, steelblue and paleturquoise have the highest correlation with traits (Fig. 2A). In order to find out the most important role in above 6 modules, we compared the module membership (MM) with gene significance (GS) in a scatter plot, which showed that genes in module orangered3 and salmon are highly correlated (Fig.S3C). GO enrichment of two modules shows that “carotene catabolic process”, “cell response to phosphate starvation”, “terpenoid biosynthesis process” and “strigolactone biosynthesis process” were the most enriched. It is also involved in plant homeostasis pathways like “trivalent inorganic anion homeostasis”, and “phosphate ion homeostasis”, as well as the biosynthesis of plant sterols and brassinosteroids (BR) (Fig.S3D). These results indicate that the genes in the modules regulate phytohormone biosynthesis and signal transduction under Pi stress.

**Figure 2.**
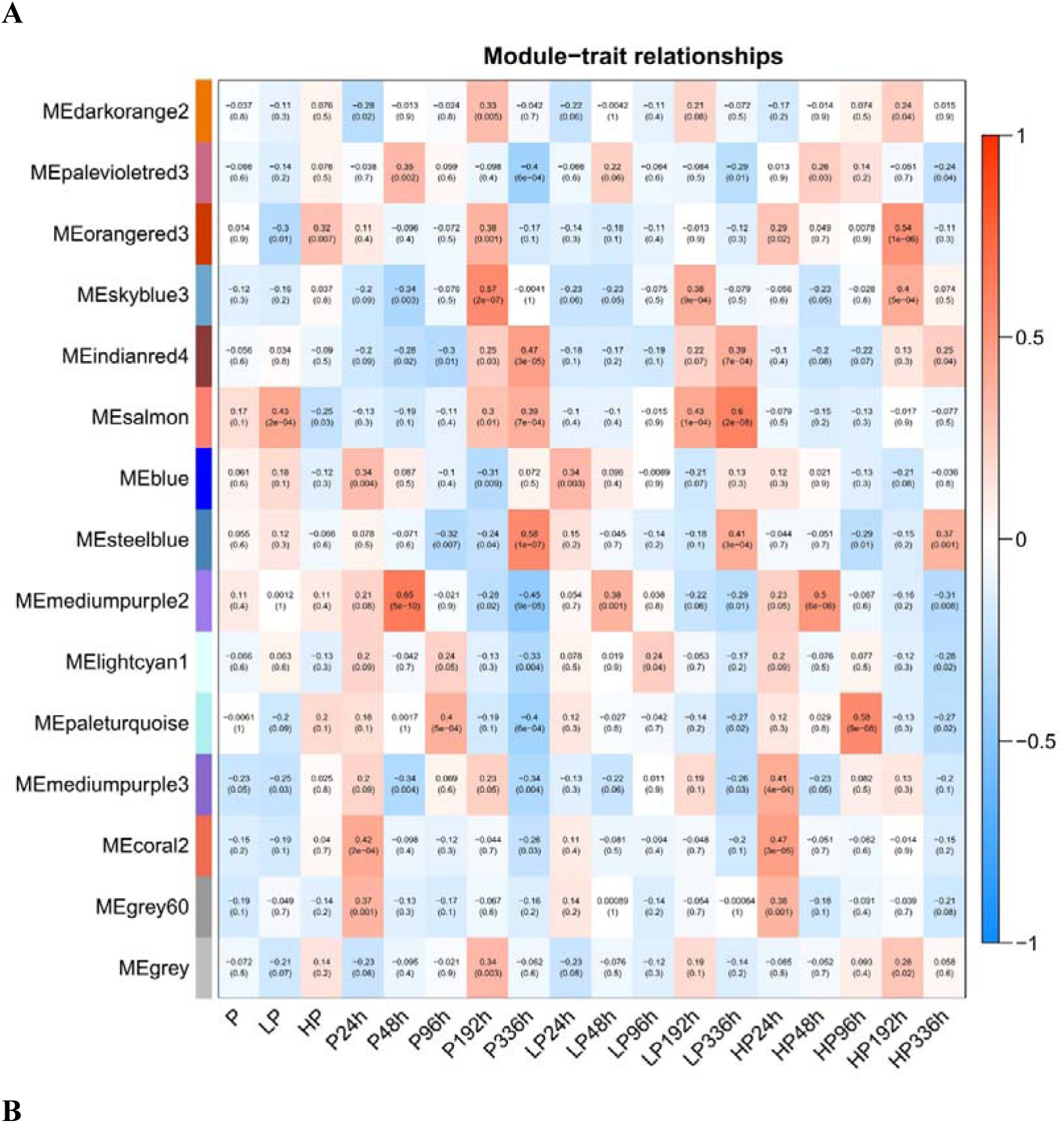
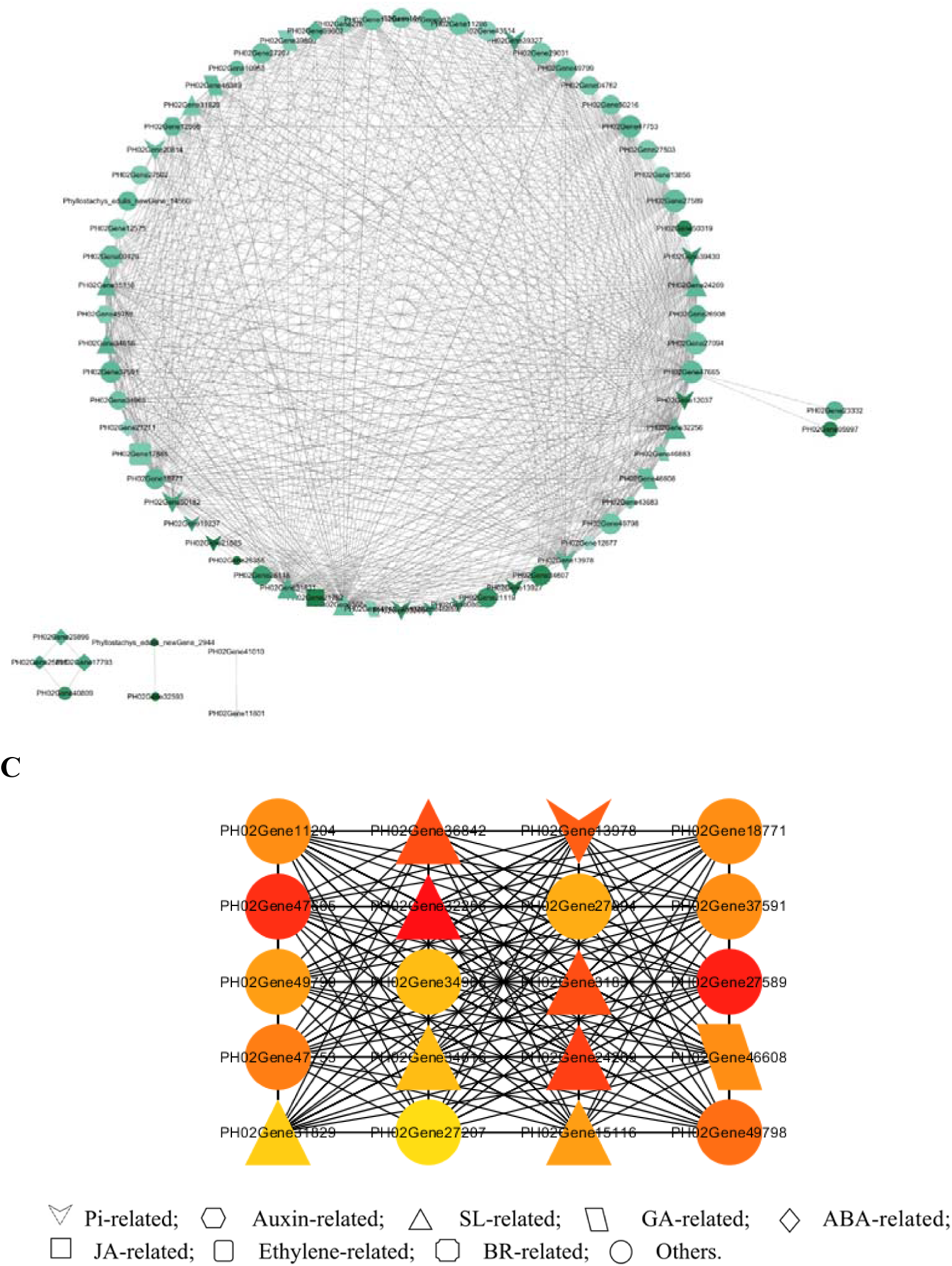
Gene co-expression network. (A) Gene co-expression network modules with traits; (B) two modules; (C) the top 20 genes with the highest connectivity. The color indicates gene significance, size represents module membership, and thickness of the lines between genes indicates the strength of the correlation.

We then constructed the co-expression network diagrams of 74 genes with weight≥0.3 in two modules and visualized them by calculating connectivity using 12 algorithms (Fig. 2B). In addition to many genes related to response to Pi starvation, there are also many biosynthesis- and signal transduction-related genes of plant hormone, including auxin, SL, gibberellin (GA), abscisic acid (ABA), jasmonic acid (JA), ethylene and brassinolide (BR). Among the 20 genes with the highest connectivity, 7 belong to SL biosynthesis and signal transduction pathway (Fig. 2C), and 4 of them ranked in the top 10%, namely *MAX1* (*MORE AXILLARY ROWTH 1*), *CCD7*(*CAROTENOID CLEAVAGE DIOXYGENASE7*), *D27* (*DWARF 27*) and *D3* (*DWARF 3*), indicating that SL pathway plays a key role in moso bamboo roots response to Pi stress. We compared these genes with the DEGs from LRP zone and primary root tips. All SL-related genes exist in DEGs most significantly enriched in the LRP zone, and several SL genes were found in primary root tips. Therefore, genes related to SL, auxin, GA, ABA, JA, ethylene and root growth among the genes significantly enriched in LRP zone and primary root tip were selected for further analysis.

### 3.5 Key candidate genes for response to Pi stress in LRP zone and primary root tip

We found 23 key genes in LRP zone and primary root tip region in response to Pi stress (Table S3). In LRP zone, there are 16 key genes including 3 genes related to PSRs and Pi transport containing SPX (Suppressor of yeast GPA1 (SYG1)/yeast phosphatase (PHO81)/human Xenotropic and Polytrophic Retrovirus Receptor1 (XPR1)) domain; 9 SL biosynthesis and signaling genes containing *MAX1*, *D27*, *CCD7*, *D10* (*DWARF10*), *D3* and *D14* (*DWARF14*); 2 auxin-related genes *ABCB1* (*ABC TRANSPORTER B FAMILY MEMBER 1*) and *NCS1* (*S-NORCOCLAURINE SYNTHASE 1*); 1 ABA-responsive gene *ERD15* (*EARLY RESPONSIVE TO DEHYDRATION 15*) and 1 cell apoptosis-related gene *ESR1* (*EMBRYO SURROUNDING REGION 1*) (Fig. 3A). Therefore, Pi-starvation responsive genes, SL biosynthesis and signaling genes and auxin-responsive genes were significantly upregulated with treatment time under low Pi with similar expression patterns. While SL receptor genes, ABA-responsive genes and cytokinin-responsive genes were significantly down-regulated under low Pi, indicating that there is complex crosstalk among hormone signals in LRP zone in response to Pi starvation.

**Figure 3.**
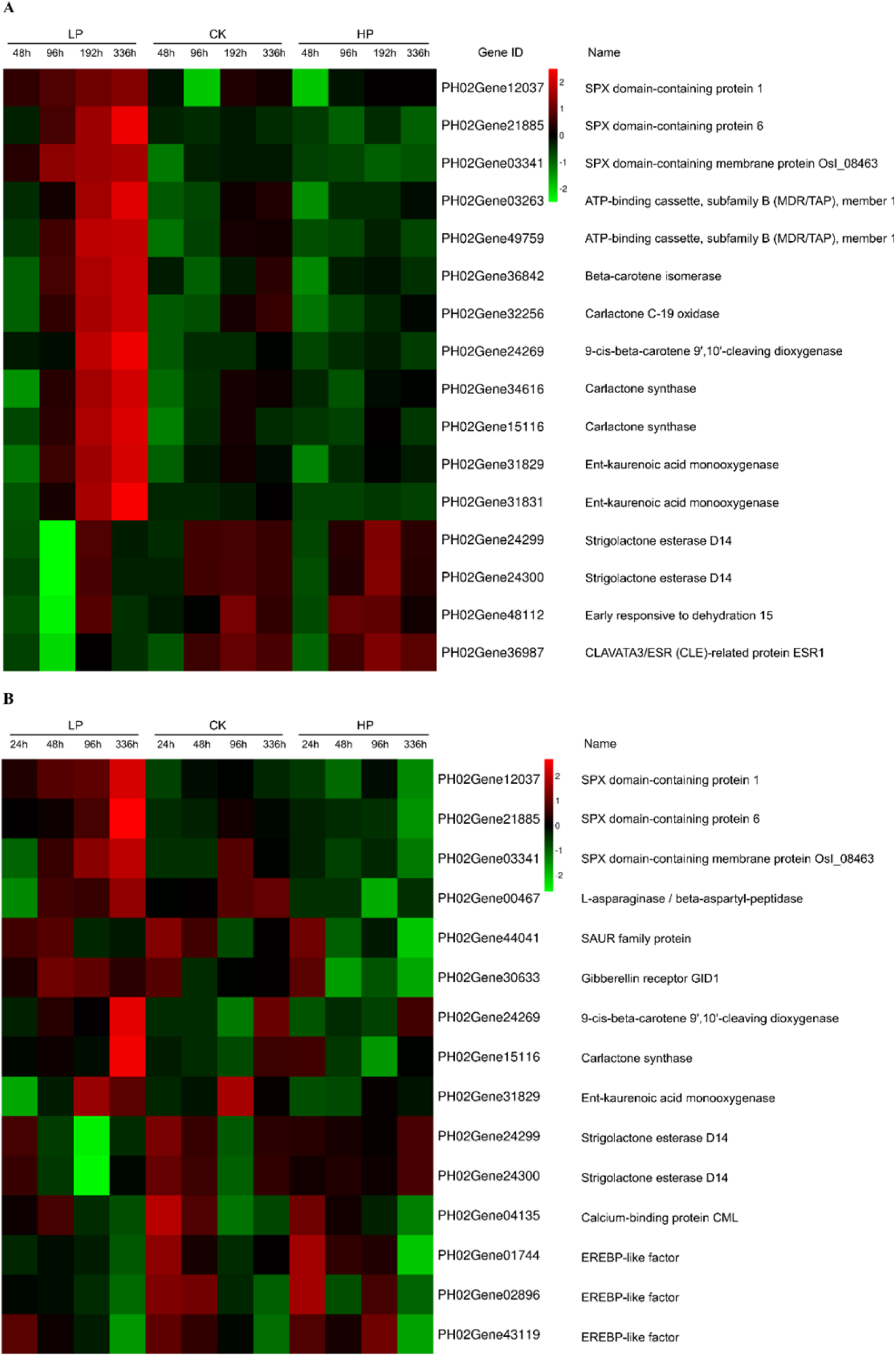
Expression trend of DEGs related to plant hormones in (A) LRP zone and (B) primary root tip of moso bamboo in response to Pi stress.

Fifteen genes were screened from the DEGs in primary root tip region including 3 Pi-responsive genes and 5 SL-related genes also found in LRP zone. Besides, there are *ASPGB1* (*ASPARAGINASE B1*) and *CML16* (*CALMODULIN-LIKE PROTEIN 16*) associated with root growth, auxin-responsive gene *SAUR11* (*SMALL AUXIN UPREGULATED RNA 11*), GA accepter *GID1* (*GIBBERELLIN INSENSITIVE DWARF1*) and 3 ethylene-responsive genes ERF4/5/071 (*ETHYLENE-RESPONSIVE TRANSCRIPTION FACTOR 4*/*5*/*071*) (Fig. 3B). Compared with the LRP zone, key genes in primary root tips did not show a continuous expression trend. However, it suggests that the primary root tip may perceive Pi signal in the environment and affect genes expression in a particular period through hormone signaling pathways to adapt to the early Pi stress. We successfully verified the expression trends of 23 key genes over time by qRT-PCR (Fig.S4, Fig.S5).

### 3.6 Role of SL in response to Pi in moso bamboo root

In order to investigate the biosynthesis and secretion of SL in bamboo roots, we measured SL content in root exudates after 96h, 192h and 336h treatment under different Pi levels, and two kinds of endogenous SL were detected. Compared with sufficient Pi, the secretion of 5-deoxystrigol (5-DS) was significantly increased under low Pi at 96h, 192h and 336h but significantly decreased under high Pi at 192h (Fig. 4A). Notably, 5-DS content at 336h under low Pi was 21.6 times and 434 times under sufficient Pi and high Pi. Regardless of Pi treatment, 5-DS content decreased gradually and significantly with time (Fig. 4B). The secretion of strigol was also significantly increased under low Pi at 192h and 336h. It slightly decreased under high Pi with no significant difference comparied with under sufficient Pi, and strigol content at 336h under low Pi was 15.2 times and 139.2 times under sufficient Pi and high Pi (Fig. 4C). Strigol content decreased gradually and significantly with time under sufficient and high Pi, but increased significantly under low Pi at 336h (Fig. 4D). These results indicate that low Pi stimulates SL biosynthesis and exudation in moso bamboo roots, and 5-DS might be the precursor of strigol in moso bamboo (Iseki et al., 2018).

**Figure 4.**
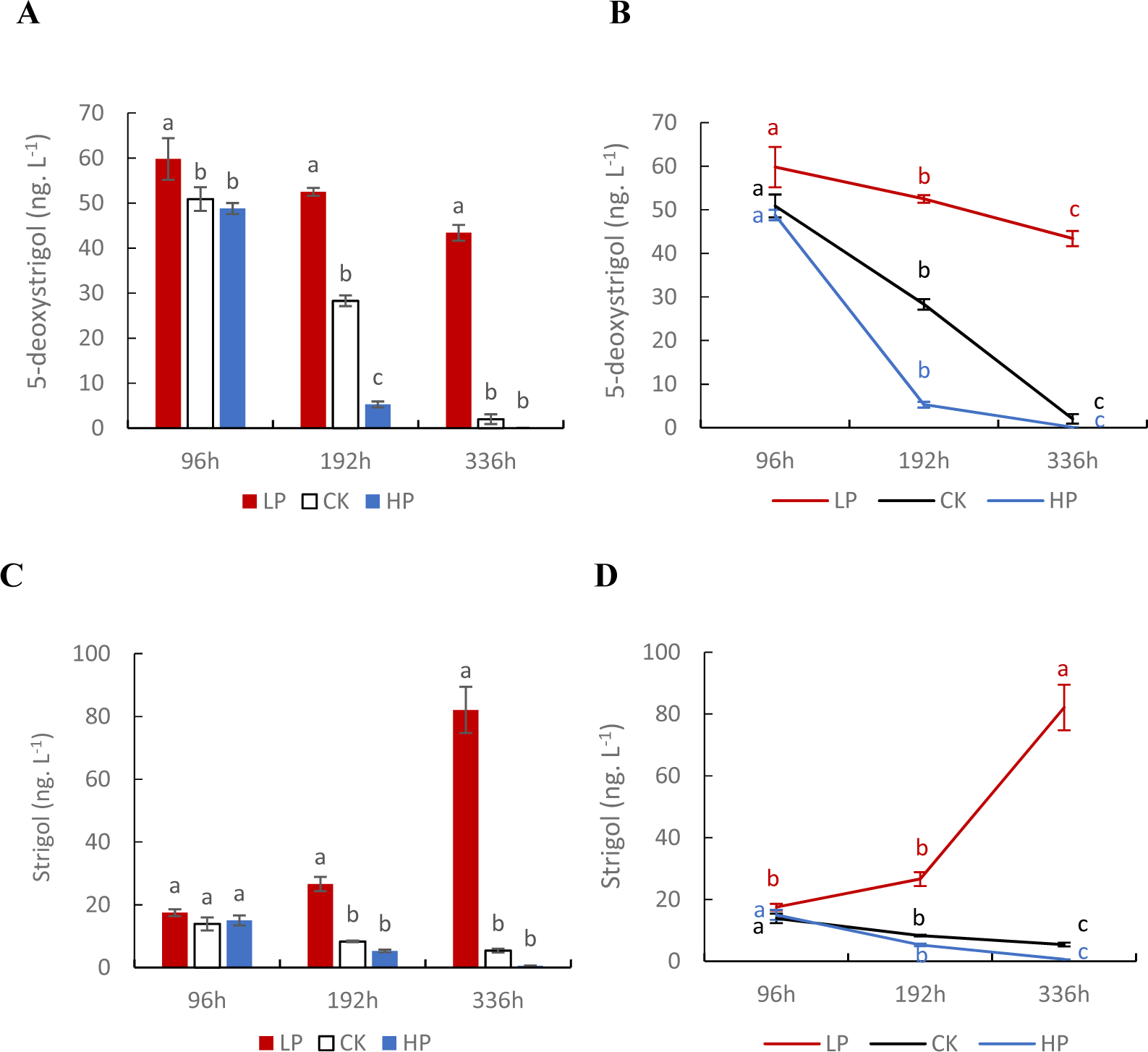
SL content in root exudates of moso bamboo at different Pi levels. Comparison of 5-deoxystrigol (5-DS) content under different Pi levels (A) and at different developmental stages (B); comparison of strigol content under different Pi levels (C) and at different developmental stages(D). (A) (C) Letters represent differences among different Pi levels at the same time, and (B) (D) differences among different developmental stages under the same Pi level.

Then we applied 0.01, 0.1, 1, 2 μM GR24 or TIS108 under sufficient Pi for 2 weeks to determine how SL regulates the growth and development of bamboo roots. With the increase of GR24 concentration, biomass decreased and root-shoot ratio had no significant difference, and 1 uM GR24 decreased both primary root length and lateral root density (Fig.S6). TIS108 concentration had no significant effect on biomass and root-shoot ratio, but 1uM TIS108 decreased primary root length and increased lateral root density (Fig.S7). To study the role of SL under Pi stress, we then conducted a compound treatment, applying 1 uM GR24 or TIS108 to low Pi, sufficient Pi and high Pi, respectively. GR24 did not change biomass at any Pi concentration but increased root-shoot ratio under low Pi, while TIS108 also had no significant effect on biomass but reduced the root-shoot ratio under low Pi (Fig.S8), suggesting that SL affects dry matter allocation of aboveground and underground parts of moso bamboo seedlings under low Pi. We also measured total phosphorus content to study the influence of SL on PAE. GR24 decreased the total phosphorus content but had no significant effect on PAE at any Pi levels, and TIS108 reduced total phosphorous content under sufficient Pi but only lowered PAE under low Pi (Fig.S9). These results show that SL promotes PAE at low Pi, but does not affect PAE when providing sufficient or excess Pi. We measured primary root length and lateral root density after treatment to analyze how GR24 and TIS108 affect root morphology under Pi stress (Fig.S10A). GR24 decreased primary root length significantly under both low and sufficient Pi but not under high Pi (Fig.S10B), indicating that GR24 can induce typical PSRs in moso bamboo, but this effect seemed to be weakened when Pi concentration was too high. TIS108 decreased primary root length significantly under all Pi levels, with the most significant reduction at low Pi (Fig.S10C), indicating that SL plays a pivotal role in the process of root growth and has a more significant impact when Pi is deficient. GR24 only decreased the lateral root density under sufficient Pi (Fig.S10D), which is also a typical PSR induced by SL in other plants like Arabidopsis. TIS108 increased the lateral root density significantly under low and sufficient Pi, but had no significant effect under high Pi (Fig.S10E), indicating that inhibition of SL biosynthesis can restore SL-inhibited lateral root formation at normal or low phosphorus concentration but not at excessive Pi concentration. Therefore, SL affects lateral root formation in a Pi-dependent manner.

### 3.7 SL affects root microscopic characteristics of root under different Pi levels

The change of root architecture under Pi stress is related to the change of anatomical structure, so we observed the microstructure of primary root tip and LRP zone under compound treatments. With the decrease of Pi concentration, the cross-section size of primary root tip increased gradually with increased cell number (Fig. 5A). This suggest that moso bamboo may adapt to environmental changes by expanding the surface area or volume of roots through cell proliferation under low Pi conditions, similar to other plants. Application of GR24 significantly reduced the cross-section size at low Pi and increased the size under sufficient and high Pi (Fig. 5A), indicating that SL affects the development of primary root tip depending on Pi concentration. TIS108 increased the size slightly under low Pi, accompanied by cell swelling and morphological disorder, but not under sufficient or high Pi (Fig. 5A). These results suggests that SL maintains cell morphological stability, especially under low Pi.

**Figure 5.**
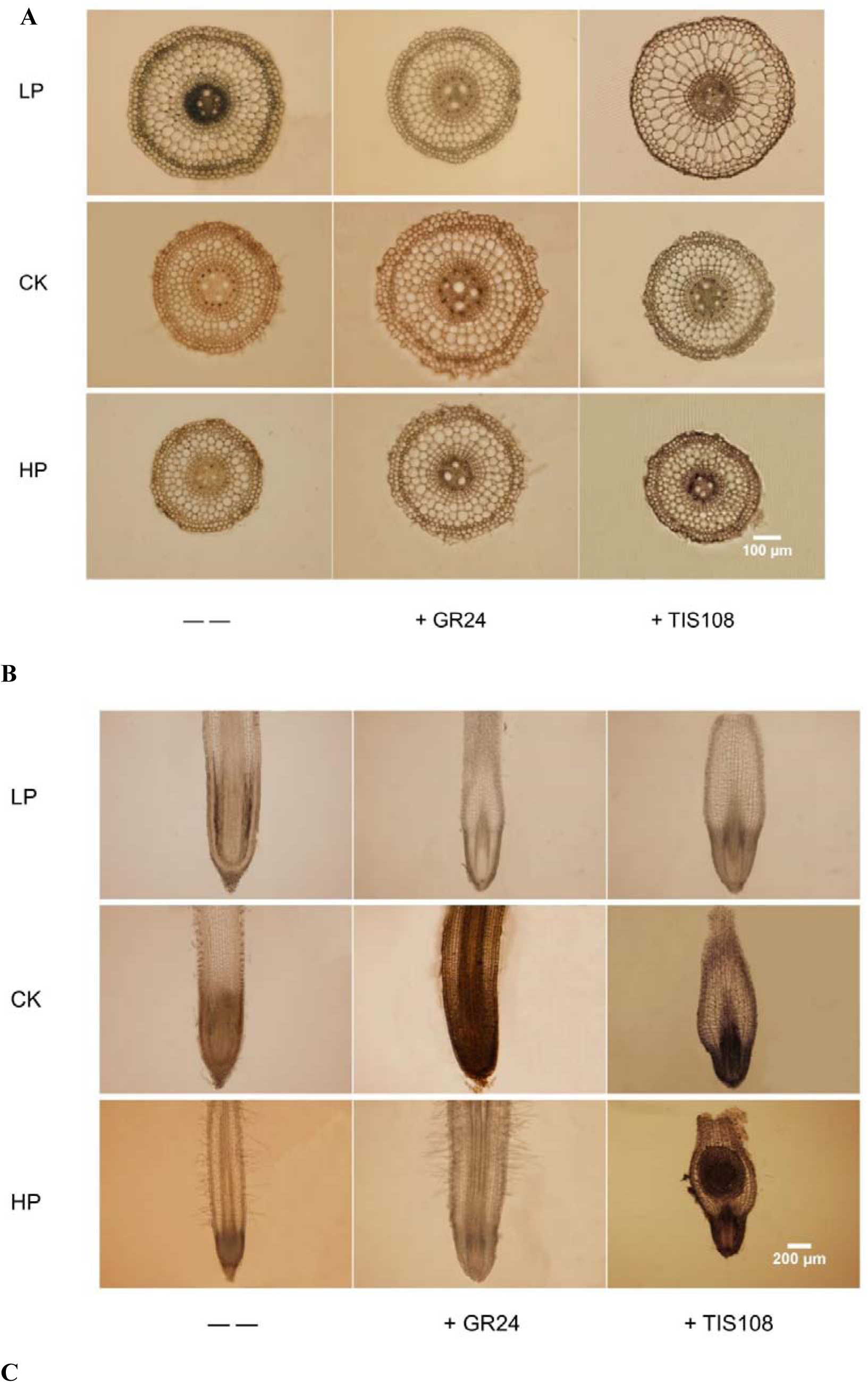
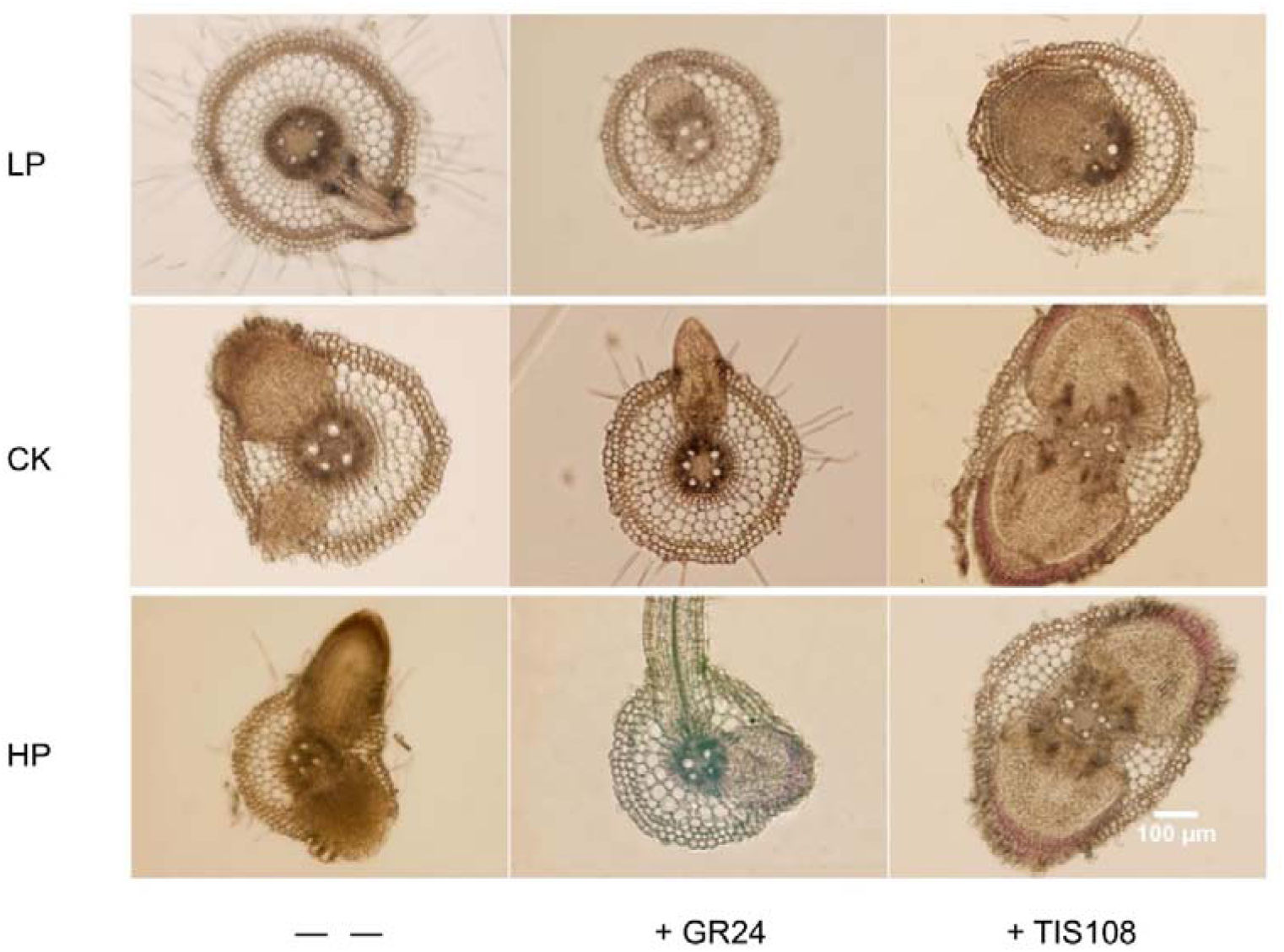
Light micrographs of moso bamboo roots treated with GR24/TIS108 at different Pi levels for 14 days. (A) transversely-sectioned primary root tip, (B) longitudinally-sectioned primary root tip, (C) lateral root formation.

The size of root apical meristem zone of primary root tip increased with the decrease of Pi level. Compared with sufficient Pi, the meristem and elongation regions of primary root tip were longitudinally longer at low Pi and shorter at high Pi. Mature region was clearly visible at high Pi at the same magnification, densely covered with root hair (Fig. 5B). GR24 reduced the size of root apical meristem and elongation zone under low Pi and slightly reduced the size under sufficient Pi. It was difficult to determine the effect of high Pi on the length of the apical meristem, but it significantly extended the distance from the root cap to the first root hair (Fig. 5B). The root cap treated by TIS108 under low Pi showed a conical shape, and the size of the root apical meristem zone was significantly shortened. Root apical meristem size was also reduced when TIS108 was applied with sufficient Pi. TIS108 application under high Pi did not affect the length of the root apical meristem but shortened the distance to the first LRP closest to the root cap (Fig. 5B). Therefore, GR24 can reduce the size of the undifferentiated root apical region at low and sufficient Pi levels but extend the size at high Pi, while SL deficiency caused by TIS108 results in shorter length of undifferentiated root apical region at low and sufficient Pi levels and affects the location of first lateral root occurrence at high Pi. These results suggest that SL plays an essential role in the process of root elongation, and this function may have a fine-tuning mechanism to regulate root elongation and lateral root formation according to different Pi concentrations.

Lateral root formation under low Pi was significantly reduced compared with that under sufficient and high Pi. There was little difference in lateral root number under sufficient and high Pi (Fig. 5C). GR24 application seemed to slow down the development of lateral root formation under low Pi and reduced the lateral root formation under sufficient Pi, but did not affect lateral root number under high Pi. TIS108 did not affect the lateral root number under any Pi levels. However, it seemed to promote more pericycle cells to form LRP, accompanied by cell stretching with a cross-section of the LRP zone became elliptical under sufficient and high Pi (Fig. 5C). These results indicate that SL affects lateral root formation and the stability of cell morphology of LRP zone in a Pi-dependent manner.

### 3.8 SL mediates response of LRP zone to Pi stress in moso bamboo

In order to investigate whether SL mediates the response of moso bamboo roots to Pi stress at the molecular level, quantitative expression analysis was conducted on 16 key genes in LRP zone under compound treatment. We compared the relative expression level of these key genes with or without GR24/TIS108 at the same time under the same Pi level.

In the LRP zone, GR24 increased *SPX1* and *SPX6* expression under sufficient and high Pi, while TIS108 decreased their expression under low Pi after 48h (Fig.S11AB). GR24 significantly increased the expression of *OsI_08463* at 96h and 192h under any Pi levels, and the increased range increased with the decrease of Pi concentration. *OsI_08463* expression dropped significantly after 192h of TIS108 treatment at low Pi (Fig.S11C). These results indicate that SL mediates the response of *SPX1*, *SPX6* and *OsI_08463* to low Pi in moso bamboo roots.

Expression of *ABCB1* and *NCS1* were significantly increased by GR24 at 96h under any Pi levels, and the increase rate was low Pi > sufficient Pi > high Pi. *ABCB1* and *NCS1* expression reached the peak at 192h after GR24 treatment with low Pi but plummeted to a level lower than that without GR24 at 336h, while the expression gradually increased after GR24 treatment with sufficient or high Pi during the whole experiment. TIS108 decreased *ABCB1* expression after 192h under low Pi but gradually increased its expression under sufficient and high Pi. TIS108 also reduced *NCS1* expression even to the level close to under sufficient Pi but slightly increased its expression under sufficient and high Pi (Fig.S11DE). These results indicate that SL mediates auxin transport in a Pi-dependent manner.

GR24 increased *MAX1* expression gradually after 96h under sufficient and high Pi, while TIS108 dramatically declined *MAX1* expression at low Pi throughout the whole period, even lower than that under high Pi (Fig.S11F). GR24 increased *D27* expression rapidly after 48h under any Pi levels, and reached the peak at 96h under low Pi, 192h under sufficient and high Pi, and then decreased gradually. TIS108 reduced *D27* expression only under low Pi after 96h to the level with high Pi (Fig.S11G). Under low Pi, *CCD7* expression was also significantly increased by GR24 at 192h but decreased at 336h, while TIS108 decreased its expression to the level at high Pi, similar to that of *MAX1* and *D27* (Fig.S11H). GR24 increased the expression levels of two *D10* at 48h and reached the peak at 192h, but the increase rate was high Pi > sufficient Pi > low Pi, while TIS108 decreased *D10* expression significantly after 96h under low Pi. Unlike expression of other SL-biosynthesis genes that were not affected by TIS108 under sufficient or high Pi, soaring expression of *D10* at 96h might cause the feedback regulation due to SL deficiency (Ito et al., 2013) (Fig.S11IJ). GR24 resulted in a significant increase in expression of two *D3* genes at 96h with the most significant jump under low Pi. *D3* expression fell after TIS108 treatment for 96h under low Pi (Fig.S11KL). Expression of two *D14* genes dipped after GR24 treatment for 48h under low and sufficient Pi but increased slightly under high Pi at 96h, while their expression was significantly increased at 192h after TIS108 treatment under low Pi (Fig.S11MN). Therefore, GR24 can improve the expression of SL-biosynthesis and -signaling genes under different Pi levels, and most genes respond earlier or increase more under low Pi. In contrast, TIS108 inhibits SL-biosynthesis genes’ expression and affects SL-signal transduction genes only under low Pi, although a few genes seem to be exceptions, like *D10*.

*ERD15* expression increased after GR24 treatment for 96h, reached the peak at 192h under sufficient and high Pi, while its expression was substantially boosted by TIS108 at 192h under any Pi levels, with the maximum value under high Pi (Fig.S11O), implying that SL mediates ABA signaling pathway in a Pi-dependent manner. GR24 had little effect or slightly reduced the expression of *ESR1* under any Pi condition, but its expression was greatly enhanced by TIS108 at 192h under high Pi (Fig.S11P), indicating that SL may affect cell proliferation by regulating *ESR1* expression, thus controlling lateral root development.

### 3.9 SL mediates response of primary root tip of moso bamboo to Pi stress

In primary root tip region, we also compared the relative expression level of 9 key genes with or without GR24/TIS108 at the same time under the same Pi level under compound treatment.

*CCD7* expression increased significantly at 48h after GR24 application with an increase rate of low Pi > sufficient Pi > high Pi, while it decreased significantly after TIS108 treatment during the whole study period only under low Pi (Fig.S12A). *D10b* expression increased significantly at 24h after GR24 treatment under low Pi and then decreased significantly to the level under sufficient and high Pi. Expression level of *D10b* decreased during the whole study period after TIS108 application under low Pi (Fig.S12B). These results indicate that GR24 can rapidly increase the expression of SL-biosynthesis genes in primary root tip of moso bamboo within 24h, and TIS108 continuously inhibited the expression of SL-biosynthesis genes, with the strongest inhibition under low Pi. *SAUR11* expression increased at 24h after GR24 treatment under low Pi and then decreased to the level at high Pi, while its expression decreased slightly during the whole study period after TIS108 application under low and sufficient Pi (Fig.S12C). These results indicate that *SAUR11* is also regulated by SL depending on Pi level when responding to auxin. *GID1* expression had no change or decreased slightly after GR24 and TIS108 treatment at different Pi levels (Fig.S12D), indicating that *GID1* in GA signaling pathway might be not regulated by SL.

*ASPGB1* expression was steadily raised after GR24 application under low and sufficient Pi but soared at 96h under high Pi and was slightly declined at 336h of three Pi levels. TIS108 reduced *ASPGB1* expression significantly after 48h under any Pi levels, with the decrease rate being low Pi > sufficient Pi > high Pi (Fig.S12E). GR24 decreased *CML16* expression in all periods under low and sufficient Pi, while TIS108 treatment increased its expression after 24h under low Pi (Fig.S12F), indicating SL might affect the primary root elongation through regulating the expression of *ASPGB1* and *CML16* under different Pi levels.

*ERF4* expression fell after GR24 treatment for 24h under sufficient and high Pi, and went up after TIS108 treatment for 24h under high Pi (Fig.S12G). GR24 reduced the expression of *ERF5* at 24h under sufficient and high Pi, while TIS108 increased its expression at 24h under low and sufficient Pi but decreased its expression under high Pi (Fig.S12H). GR24 increased *ERF071* expression after 48h and decreased at 336h under low and sufficient Pi, while TIS108 reduced its expression at 24h under low Pi (Fig.S12I). These results suggest that SL mediates the expression of *ERF4*, *ERF5* and *ERF071* in a Pi-dependent manner.

### 3.10 SL-mediated gene regulatory network response to Pi stress in moso bamboo

Through transcriptome sequencing and relative expression analysis of 23 key genes under different Pi levels and SL-analog or -inhibitor treatment, we obtained the gene regulatory network of SL mediated LRP zone and primary root tip in response to Pi stress (Fig. 6). In primary root tip, SL inhibits the expression of *CML16*, *ERF4*, *SPXs* under low Pi to promote root elongation (Fig. 6A). In contrast, the loss of SL under high Pi relieves the inhibition of *ERF071* and *ERF5* expression and promotes *ASPGB1* expression, thus inhibiting root elongation (Fig. 6B). In LRP zone, SL reduces lateral root formation by promoting *SPXs*, *ABCB1* and *NCS1* while inhibiting the expression of *ESR1* and *ERD15* under low Pi (Fig. 6A). Under high Pi, the absence of SL inhibits the expression of *SPXs* and releases the promotion of *ABCB1* and *NCS1* expression and the inhibition of *ESR1* and *ERD15* expression, leading to increased lateral root formation (Fig. 6B).

**Figure 6.**
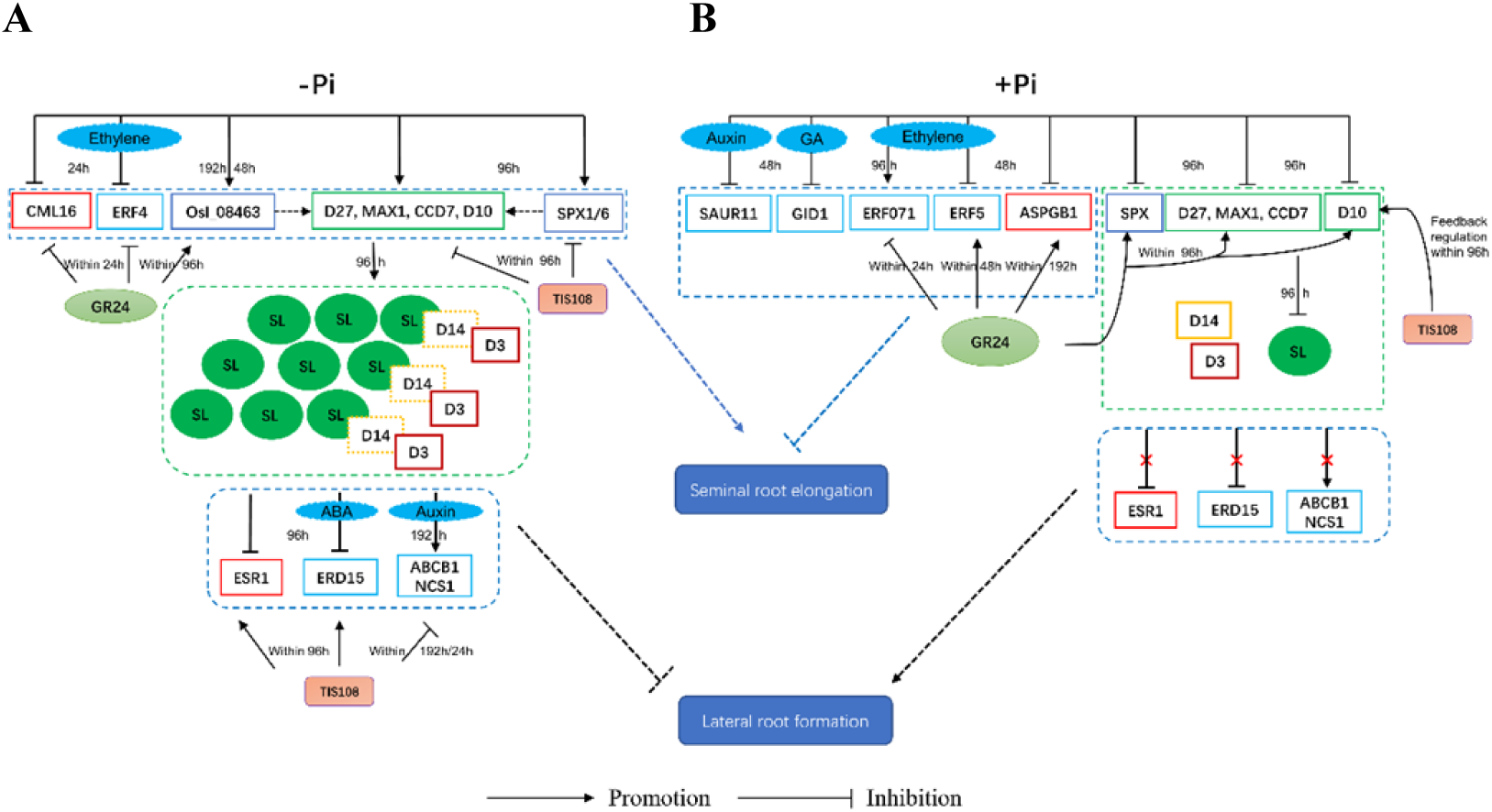
SL-mediated gene regulatory network of LRP zone and primary root tip in response to Pi stress in moso bamboo.

## 4. Discussion

### 4.1 SL affects root development of moso bamboo under Pi stress

Lack of available Pi in soil is a global problem and a significant limiting factor for agricultural production and crop yield improvement. Our results show that the root architecture of moso bamboo varies under different Pi levels. Primary root elongation is promoted, and lateral root density is reduced under low Pi, similar to that of rice (Sun et al., 2014); while primary root length is inhibited and lateral root formation is promoted, similar to that of Arabidopsis (Linkohr et al., 2002), suggesting that moso bamboo may have its own unique molecular response mechanism. The adaptive mechanism of phytohormone regulation of root growth under Pi stress is quite complex and mainly depends on their interaction. Many studies have shown that SL participates in low Pi regulation of plant root growth and development. Pi level in plants is correlated with SL exudation, and Pi deficiency can promote the biosynthesis and secretion of SL (Yoneyama et al., 2012). GR24 can induce PSRs, such as root hair elongation, anthocyanin accumulation, acid phosphatase production and plant weight reduction (Ito et al., 2015), suggesting that there may be an overlap between these two signals and plant homeostasis pathways. By applying GR24 and TIS108 to different Pi levels, we investigated how SL affected the growth and development of bamboo roots depending on Pi levels. During the 14-day study period, GR24 significantly increased and TIS108 decreased root-shoot ratio under low Pi, indicating that SL’s influence on biomass allocation is dependent on Pi concentration. GR24 reduced primary root length under both low and sufficient Pi similar to that of Arabidopsis but as opposed to that of rice, and also reduced lateral root density under sufficient Pi similar to that of rice but opposite to Arabidopsis (Sun et al., 2014; Marzec et al., 2018), indicating that SL may result in the unique mechanism of moso bamboo roots in response to Pi stress. TIS108 inhibited primary root growth of moso bamboo at any Pi levels like many plants, suggesting that SL plays an important role in root elongation even in the case of excess Pi. TIS108 and enhanced lateral root density only under low and sufficient Pi, indicating that SL controls the occurrence of lateral roots depending on Pi level in moso bamboo.

### 4.2 SL regulates expression of Pi homeostasis-related genes

There have been many studies on the genes of low Pi stress. When available Pi is scarce in soil, plants will sense low Pi signal and go through a series of signal transduction processes to regulate the expression of Pi-related regulatory and structural genes. In Arabidopsis and rice, many Pi starvation responsive genes are controlled by PHR1 (PHOSPHATE STARVATION RESPONSE REGULATOR 1), but the expression of *AtPHR1* and *OsPHR2* does not respond significantly to Pi starvation. Instead, plants regulate PHR1 by sensing changes in Pi level in cells to activate the central regulator SPXs (Wang et al., 2014). SPX1 and SPX2 are Pi-dependent inhibitors of PSR1, and SPX1/PHR1 module connects Pi sensing and signal transduction (Puga et al., 2014); and SPX6 negatively regulates Pi starvation by inhibiting the transcription factor PHR2 (Zhong et al., 2018). In legumes, *PvSPX1* overexpression resulted in increased Pi concentration in the root, inhibited root growth, and enlarged root hair area in transgenic plants (Yao et al., 2014). There are few studies on the effects and molecular mechanisms of excessive Pi stress on plants, but low-affinity and high-affinity Pi transporter systems have been found in plants. High-affinity Pi transporters are transcriptionally induced in the micromole range at low Pi concentrations, while genes for low-affinity Pi transporters are constitutively expressed and function in the nanomole range at relatively high Pi concentrations. For example, *PtPHT4;1a* in poplar (*Populus trichocarpa*) was up-regulated under low Pi, while *PtPHT4;1b* and *PtPHT4;5b* were up-regulated under high Pi, suggesting that they play different roles in response to Pi supply (Zhang et al., 2016).

SL-regulated signal transduction is also an important part of plant adaptation to Pi stress. SL can regulate the activity of a series of Pi starvation-related genes to improve Pi signal transduction, thus improving PAE and regulating root architecture (Kumar et al., 2015). PAE in wheat is related to the regulation of PHO2 activity and enhancement of Pi signaling, and PHO2 activity is regulated by SL (de Souza Campos et al., 2019). In tomatoes, Pi deficiency results in strongly upregulation of SL yield and expression of PSR genes such as *PHT* and *SPXs*, acid phosphatase secretion, and increased root shoot ratio (Wang et al., 2021b). Under Pi-limited conditions, *SPX1* and *SPX3* in alfalfa (*Medicago truncatula*) promote the expression of SL biosynthesis gene *D27*, increasing arbuscular mycorrhizal (AM) branching in response to root exudates, and *SPX1* and *SPX3* redundantly control arbuscular degradation in AM cells (Wang et al., 2021a). The target genes of *PHR2* detected by ChIP-seq and RNA-seq included SL-biosynthesis gene *CCD7*, and SL release reduced in the exudate of *phr2* mutant (Das et al., 2021).

Our results show that key DEGs screened from the LRP zone mainly exist between low Pi and sufficient/high Pi, and the AFC of DEGs between high Pi and low Pi is slightly higher than those between sufficient Pi and low Pi, suggesting that the molecular mechanism response to low Pi stress may be more complex. In Arabidopsis, SL regulates lateral root formation under Pi deficiency conditions through inhibiting auxin transport and increasing auxin perception by PIN1/2 and TIR1, respectively (Koltai, 2015; de Souza Campos et al., 2019). However, no auxin-signaling gene *TIR1* or another auxin transport gene *PIN* was found. DEGs significantly enriched in primary root tips are also located between low Pi and sufficient/high Pi with a scattered expression trend. Expression patterns of *SPX1*, *SPX6*, *OsI_08463* and *CCD7*, *D10*, *D3*, *ASPGB1* are similar, but *D27*, *CCD8*/*D10*, *MAX1* are not detected as DEGs as in LRP zone. This may be related to tissue specificity and sampling time between primary root tip and LRP zone. When GR24 was applied to LRP zone of moso bamboo under different Pi concentrations, Pi transporter genes *OsI_08463*, SL-biosynthesis genes *CCD7*, *D27*, *MAX1*, SL-signaling genes *D3*, and auxin transporter genes *ABCB1* and *NCS1* were upregulated from 48h to 192h, and the increase rate was the highest under low Pi. *SPX1*, *SPX6* and *D10* were also regulated at 24h and 48h, but the increase rate was greater under sufficient or high Pi. Expression of most genes returned to the level without GR24 at 336h, indicating the discontinuity of GR24 treatment effect or some other regulatory mechanism was caused. All SL-biosynthesis and transduction genes were downregulated rapidly and significantly when TIS108 was applied under low Pi, but not under sufficient or high Pi. *CCD7* and *D10* were downregulated when GR24 was applied to primary root tips of moso bamboo under different Pi levels, with the most significant decrease under low Pi. These results indicate that SL can regulate Pi deficiency responsive genes, and exogenous GR24 can induce PSRs under sufficient and high Pi conditions.

## 5. Conclusion

In summary, we investigated the root morphology of moso bamboo under different Pi levels at physiological, cellular and molecular levels. We explored the interaction between Pi signal and homeostasis and SL regulation in root elongation and branching, and the mechanism of SL mediating the response of moso roots to Pi stress. As Pi concentration decreases, primary root elongates gradually with extended root apical meristem, and lateral root density decreases with reduced LRP. Root-shoot ratio increased gradually, and endogenous SL secreted by roots increased significantly. SL pathway plays a key role in response of root to Pi stress, especially in low Pi conditions, by influencing PAE and altering the proportion of biomass to tissues. Pi starvation responsive genes have strong interactions with SL-related genes. SL affects the primary root elongation and branching by regulating its biosynthesis and signal transduction. This process also has complex interactions with auxin, ABA, ethylene and other hormone signals.

## Acknowledgements

This work was funded by grants from the National Natural Science Foundation of China (grant numbers 31870656) and the Zhejiang Provincial Natural Science Foundation of China (grant number LZ19C160001).

## Conflict of interests statement

The authors declare that they have no conflict of interest.

## Author contributions

Q.W. collected plant materials, analyzed and interpreted the data, and wrote the manuscript. P.Y. and M.A. revised the manuscript. RY.G. supervised the data analyses. MB.Z. designed the research and revised the manuscript. All authors read and approved the final manuscript.

## Figure legends

Figure S1 Effect of Pi availability on moso bamboo seedlings grown for 4 weeks in hydroponic solution with different Pi concentrations.

Figure S2 Effect of Pi stress on root morphology of moso bamboo for 4 weeks.

Figure S3 WGCNA analysis of all gene in primary root tip region and LRP zone of moso bamboo.

Figure S4 Comparison of relative expression levels between qRT-PCR and transcriptome of sixteen key genes in LRP zone.

Figure S5 Comparison of relative expression levels between qRT-PCR and transcriptome of nine key genes in primary root tips.

Figure S6 Effects of GR24 on (A) biomass, (B) root-shoot ratio, (C) primary root length and (D) lateral root density of moso bamboo.

Figure S7 Effects of TIS108 on (A) biomass, (B) root-shoot ratio, (C) primary root length and (D) lateral root density of moso bamboo.

Figure S8 Effect of GR24/TIS108 on biomass and root-shoot ratio of moso bamboo under Pi stress.

Figure S9 Effect of GR24/TIS108 on total phosphorous content and phosphorus acquisition efficiency (PAE) of moso bamboo under Pi stress.

Figure S10 Effect of GR24/TIS108 on primary root length and lateral root density of moso bamboo under Pi stress.

Figure S11 Relative expression of sixteen key genes in LRP zone of moso bamboo plants treated with GR24/TIS108 under different Pi levels.

Figure S12 Relative expression of nine key genes in primary root tips of moso bamboo plants treated with GR24/TIS108 under different Pi levels.

Table S1 Significantly enriched gene ontology (GO) terms and items in LRP zone of moso bamboo in response to Pi stress.

Table S2 Significantly enriched gene ontology (GO) terms and items in primary root tips of moso bamboo in response to Pi stress.

Table S3 Annotation and function of key genes in LRP zone and primary root tips of moso bamboo in response to Pi stress.

Table S4 Genes and primer sets used for qRT-PCR.

